# Synaptic diversity naturally arises from neural decoding of heterogeneous populations

**DOI:** 10.1101/2020.10.15.341131

**Authors:** Jacob L. Yates, Benjamin Scholl

## Abstract

The synaptic inputs to single cortical neurons exhibit substantial diversity in their sensory-driven activity. What this diversity reflects is unclear, and appears counter-productive in generating selective somatic responses to specific stimuli. We propose that synaptic diversity arises because neurons decode information from upstream populations. Focusing on a single sensory variable, orientation, we construct a probabilistic decoder that estimates the stimulus orientation from the responses of a realistic, hypothetical input population of neurons. We provide a straightforward mapping from the decoder weights to real excitatory synapses, and find that optimal decoding requires diverse input weights. Analytically derived weights exhibit diversity whenever upstream input populations consist of noisy, correlated, and heterogeneous neurons, as is typically found *in vivo*. In fact, in silico weight diversity was necessary to accurately decode orientation and matched the functional heterogeneity of dendritic spines imaged in vivo. Our results indicate that synaptic diversity is a necessary component of information transmission and reframes studies of connectivity through the lens of probabilistic population codes. These results suggest that the mapping from synaptic inputs to somatic selectivity may not be directly interpretable without considering input covariance and highlights the importance of population codes in pursuit of the cortical connectome.

## Introduction

Cortical neurons are driven by large populations of excitatory synaptic inputs. Synaptic populations ultimately shape how sensory signals are encoded, decoded, or transformed. The sensory representation or functional properties of an excitatory input population will define and constrain the operations a neuron can perform and reflects the rules neurons use to form connections. Electrophysiological and anatomical studies suggest that connections between excitatory neurons exhibits functional specificity, where inputs are tuned for similar features as the soma (Cossell et al., 2015; Ko et al., 2011; Lee et al., 2016; Reid and Alonso, 1995). In contrast, synaptic imaging techniques have revealed that synaptic populations exhibit functional diversity, deviating from canonical connectivity rules, such as ‘like-connects-to-like’ (Scholl and Fitzpatrick, 2020). This functional diversity within input populations has been observed in a variety of mammalian species, from rodents to primates, and for a variety of sensory cortical areas (Chen et al., 2013, 2011; Iacaruso et al., 2017; Jia et al., 2011, 2010; Ju et al., 2020; Kerlin et al., 2019; Scholl et al., 2017; Wertz et al., 2015; Wilson et al., 2018, 2016). This apparent discrepancy challenges our understanding about how synaptic inputs drive the selective outputs of cortical neurons and leads to a simple fundamental question: If the goal is to produce selective somatic responses, why would a neuron have excitatory synaptic inputs tuned far away from the somatic preference?

To answer this question, we turn to population coding theory; starting with the idea that to accurately represent sensory signals, cortical neurons must decode the activity of upstream populations. This decoding is likely accomplished by combing signals across neural populations (Graf et al., 2011; Jazayeri and Movshon, 2006). Many studies have examined how sensory variables might be decoded from cortical populations (Butts and Goldman, 2006; Graf et al., 2011; Shamir and Sompolinsky, 2006), an endeavor increasingly applied to larger population sizes with innovative recording techniques (Rumyantsev et al., 2020; Stringer et al., 2019). These decoding approaches are often used as a tool to quantify the information about a stimulus available in a neural population, carrying the assumption that downstream areas could perform such a process (Berens et al., 2011; DiCarlo et al., 2012). In real brain circuits, decoders must be composed of individual neurons, driven by sets of synaptic inputs, akin to a decoder’s weights over a given input population. To date, few studies have explicitly examined the weight structure of population decoders (Jazayeri and Movshon, 2006; Rust et al., 2006; Zavitz and Price, 2019).

In this paper, we investigate the weights of a simple population decoder and how they compare to real synaptic inputs measured *in vivo*. Focusing on a single sensory variable, orientation, we derive the maximum-likelihood readout for a simulated input population that encodes stimuli with noisy tuning curves (e.g., Ecker et al., 2011). Under reasonable assumptions, the decoder weights can be interpreted as the synaptic connectivity between the input population and the downstream decoder neurons. This allows us to examine how synaptic connectivity depends on properties of the input population and to directly compare population decoders to synaptic input measured *in vivo*. We then test a hypothesis that an optimal decoder will show substantial heterogeneity in its synaptic weights given a biologically realistic input population. We find that when input populations are shifted copies of the same tuning curve, the synaptic excitatory inputs closely resemble the somatic output. However, with a biologically realistic input population, the expected inputs onto readout neurons exhibit functional diversity. We then compare the orientation tuning of simulated inputs with large populations of dendritic spines (excitatory synaptic inputs) onto individual neurons of ferret primary visual cortex (V1), recorded with two-photon calcium imaging *in vivo*. This comparison revealed similar diversity in the orientation tuning of dendritic spines on ferret V1 neurons and simulated decoder weights. The similarity between the synaptic populations of actual V1 neurons and the optimal neural decoder suggest that diversity and heterogeneity observed in dendritic spines across sensory cortices are, in fact, expected when considering how information is propagated through neural circuits in the presence of noise.

## Results

Following several decades of work on population coding theory, we derive a Bayesian decoder to report the probability of a visual stimulus given inputs from a neural population (Fig. 1). With this framework, and given the specifics of the *encoding* population, we can analytically derive the optimal *decoding* weights of a population of readout neurons. Here, we use “optimal” to refer to the maximum-likelihood solution. Previous work has shown that a population of neurons could perform such probabilistic decoding with weighted summation and divisive normalization, as long as their inputs exhibit Poisson-like noise (Jazayeri and Movshon, 2006; Ma et al., 2006). Starting from that basic framework, we derived a decoder that represents the probability that each possible stimulus orientation was present given the responses of a large population of upstream, input neurons (*P_IN_*). This is effectively a categorical decoder, where each possible orientation is a different category. Similar decoders have been used throughout the literature to estimate how much information is in a neural recording and suggest how downstream neurons might read it out (Graf et al., 2011; Stringer et al., 2019). Our decoder has weight vectors for each possible stimulus orientation, which integrate across *P_IN_* and are passed through a static nonlinearity (the exponential function) and normalized. As we will show below, given specific assumptions about the variability in *P_IN_*, the weights over *P_IN_* depend systematically on the tuning functions and covariance of *P_IN_*. Following a characterization of this decoding framework, we will make direct comparisons with real data: defining an effective “synaptic input population” (*P_SYN_*) as nonzero, positive weights over *P_IN_*. Although our strategy applies to any one-dimensional stimulus variable, we describe this model in the context of orientation of drifting gratings presented to V1 neurons for a direct comparison with *in vivo* measurements.

**Figure 1:**
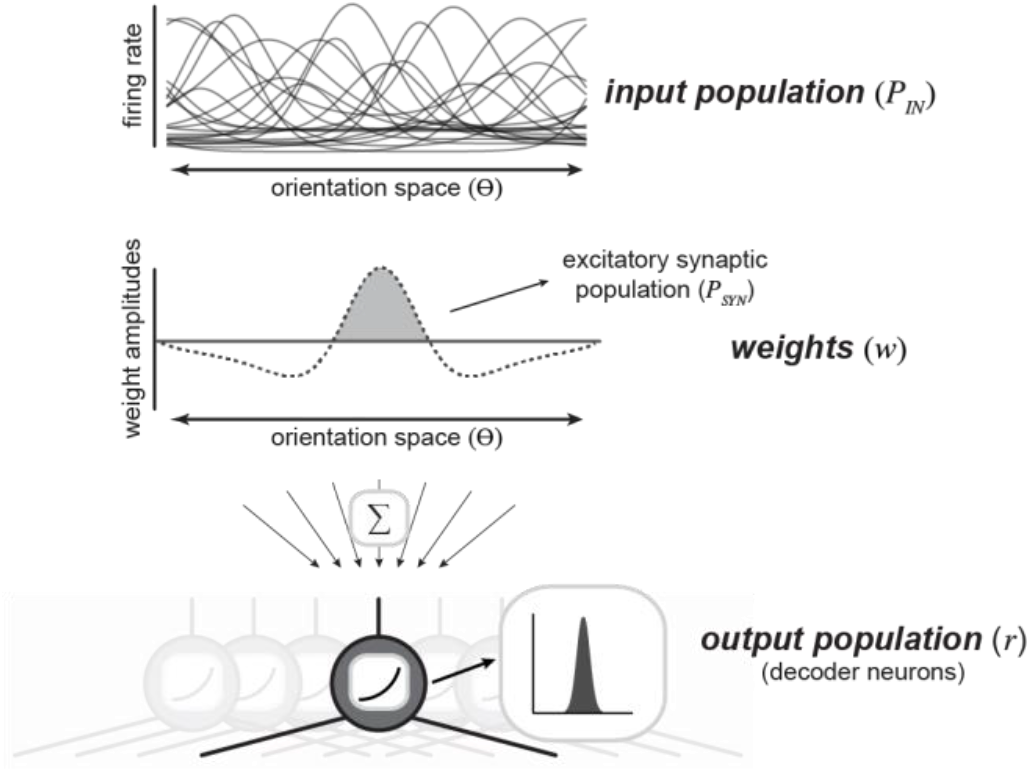
A population decoding framework to study synaptic diversity. An upstream population of neurons are tuned for a single stimulus variable (orientation) (*top*). This input population is readout by downstream decoder neurons (*bottom*). Downstream neurons decode stimulus identify by reading out spikes from the upstream input population. Each decoder neuron is defined by a set weights (*middle*) over the upstream population, which are summed and rectified to produce an output.

### A neural population as a probabilistic decoder

A categorical probabilistic decoder reports the probability that a particular stimulus orientation, θ*_k_*, was present given the spiking responses of an input population, ***R***. This can be expressed as a normalized exponential function of the log-likelihood plus the log prior for each θ*_k_*,

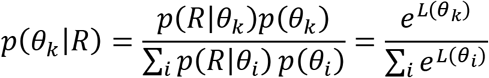

where

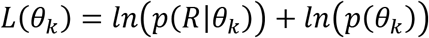

The likelihood, *p*(***R*** | θ*_k_*), is the probability of the observed responses in an input population given the stimulus *k* and *p* (θ*_k_*) is the prior probability of that stimulus class. If *p* (***R*** | θ*_k_*) is in the exponential family, then *L* (θ*_k_*) can be written as a weighted sum of the input population response vector plus an offset, which can be estimated numerically via multinomial logistic regression (Ma et al., 2006). For simplicity, we assume the input population has a response that is a function of the stimulus plus Gaussian noise, and equal covariance across all stimulus conditions. Although this assumption about the covariance structure deviates from real neural activity, this assumption means the weights and offset can be solved analytically (see Methods), and as will be shown below, such a simple model makes substantial headway in explaining biological phenomena. Our goal here is to provide a plausible alternative to “somatic selectivity” for the connectivity rules in cortex. Under the Gaussian assumption, the decoder amounts to:

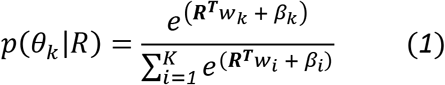

where

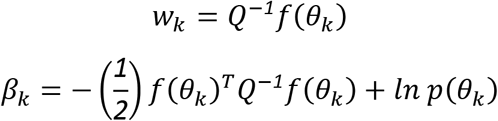

Here, *f* (θ*_k_*) is the mean input population response to stimulus orientation, *k, K* is the total number of orientations, and *Q* is the covariance matrix. The covariance term captures the influence of each neuron’s response variance (diagonal elements) and the variability shared with other neurons (off-diagonal elements). Intuitively, in the absence of covariability (i.e., off-diagonal elements are zero), the weights are proportional to the signal-to-noise ratio of the neuron (the mean divided by the variance). The term ***R^T^****w_k_* is the dot product between the population response and weights. The second term, *β_k_* is an offset for each stimulus. *ln p*(*θ_k_*) is a constant reflecting the log prior probability of stimulus *k*. It is worth noting that if the covariance depends on the stimulus, the optimal readout is no longer a linear function of ***R*** and is quadratic, which can be interpreted as a complex-cell (Jaini and Burge, 2017; Pagan et al., 2016) and is a potentially fruitful future direction.

In this study, we focus on the weights of this simple Gaussian, equal covariance decoder in order to examine how synaptic tuning from such a simple decoder would arise. Because the optimal weights have an analytic solution (eq. 1), we can see how they depend on the parameters of *P_IN_*. The simplifying assumptions we use to derive the maximum-likelihood weights help build intuitions about what can be expected in biological circuits, and linear weights such as these could be learned by real neural systems (Dayan and Abbott, 2001). A key difference here from prior work is that rather than focus on discrimination (Haefner et al., 2013), we treat orientation estimation as a multiclass identification problem, discretizing *θ* such that for each possible θ*_k_*, there is a separate weight vector. Thus, in this derivation, the optimal weights depend on the tuning curves themselves, not the derivative.

### Characteristics of a neural population decoder

To understand how synaptic weights depend on input statistics, we derived maximum-likelihood weights for input populations, *P_IN_*, with different tuning and covariance. To generate *P_IN_* we simulated *N* neurons responding to *K* oriented stimuli (θ = [−90°:*K*/180:+90°]). We briefly describe the construction of *P_IN_* here (full details are described in the Methods). Each neuron is defined by a tuning function and noise term, describing trial-by-trial variability, which are summed to generate stimulus-driven responses. We compared two fundamentally different types of input populations that have been used in the literature, homogeneous and heterogenous, as well as the role of correlated variability in shaping readout weights. A homogeneous *P_IN_* consists of shifted copies of a single tuning curve (Fig. 2a). Heterogenous *P_IN_* have diverse tuning functions and were generated to match measurements from macaque V1 (Ringach et al., 2002). The heterogenous *P_IN_* consisted of tuning curves closely resembling V1 physiology in terms of the variation in peak firing rate, bandwidth, and baseline firing rate (Fig. 2d). Varying amounts of limited-range correlations were included such that the noise correlation between two neurons depends on the difference in their tuning preferences (Ecker et al., 2011; Kohn et al., 2016).

**Figure 2:**
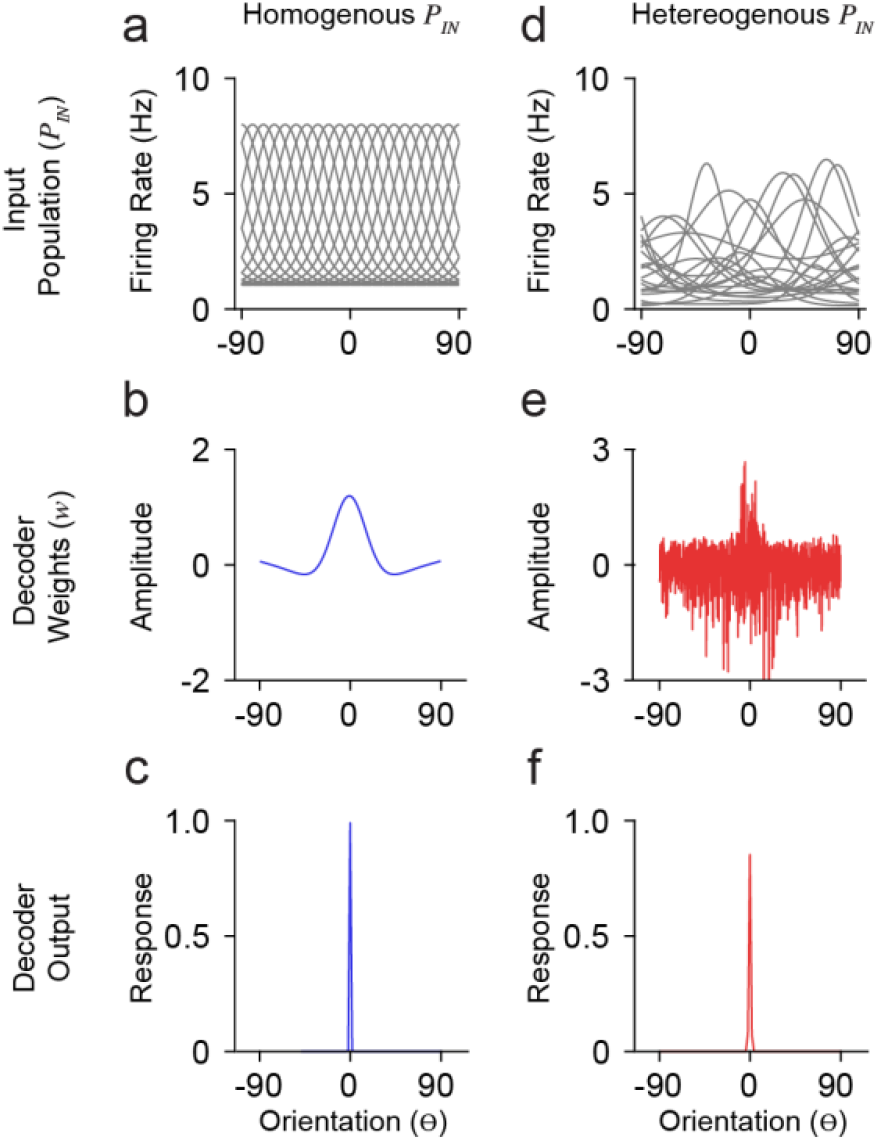
Model simulations with homogenous and heterogeneous input populations. (a) Orientation tuning of a homogenous input population. Shown is a subset of the total population (n = 20/1000). Ordinate is orientation preference, restricted between −90° and 90° (b) Derived weights for a single decoder neuron (preferring 0°) reading out the homogenous (*blue*) input population in (a). Weights for homogenous populations smoothly vary over orientation space. (c) Response output of the decoder neuron whose weights are shown in (b). (d-f) Same as in (a-c) for a heterogeneous input population with moderate correlation (*c_o_* = 0.25). Note that decoder weights for heterogeneous input populations are not smooth.

The statistics of *P_IN_* responses, ***R***, will dictate the weight structure for neurons in a decoding population. For a homogeneous *P_IN_*, the weights are smooth across orientation space and exhibit three primary features: a prominent peak about the preferred orientation of the output tuning, slight negative weights for orientations just outside the preferred, and near-zero weights at orthogonal orientations (Fig. 2c). With more realistic tuning diversity (heterogenous *P_IN_*), optimal weights are no longer smooth (Fig. 2d-e). While the optimal weights appear to roughly have the same overall shape as for homogeneous *P_IN_*, there is considerable positive and negative weighting across orientation space. Despite substantial changes in optimal weight vectors, the decoder output (i.e. somatic response) tuning was narrow (Fig. 2f), similar to the output for the homogeneous case (Fig. 2e).

To explore the importance of decoding weight diversity, we imposed a smoothing penalty on weight vectors (Park and Pillow, 2011). We calculated cross-validated decoder accuracy using the mean-squared error between the maximum a posteriori estimation and true stimulus (see Methods). Different degrees of smoothing are shown for an example set of weights in Figure 3a. We simulated a range of population sizes (*N* = 2 - 2048) and correlations (*c_o_* = 0, 0.25, 0.50).

**Figure 3:**
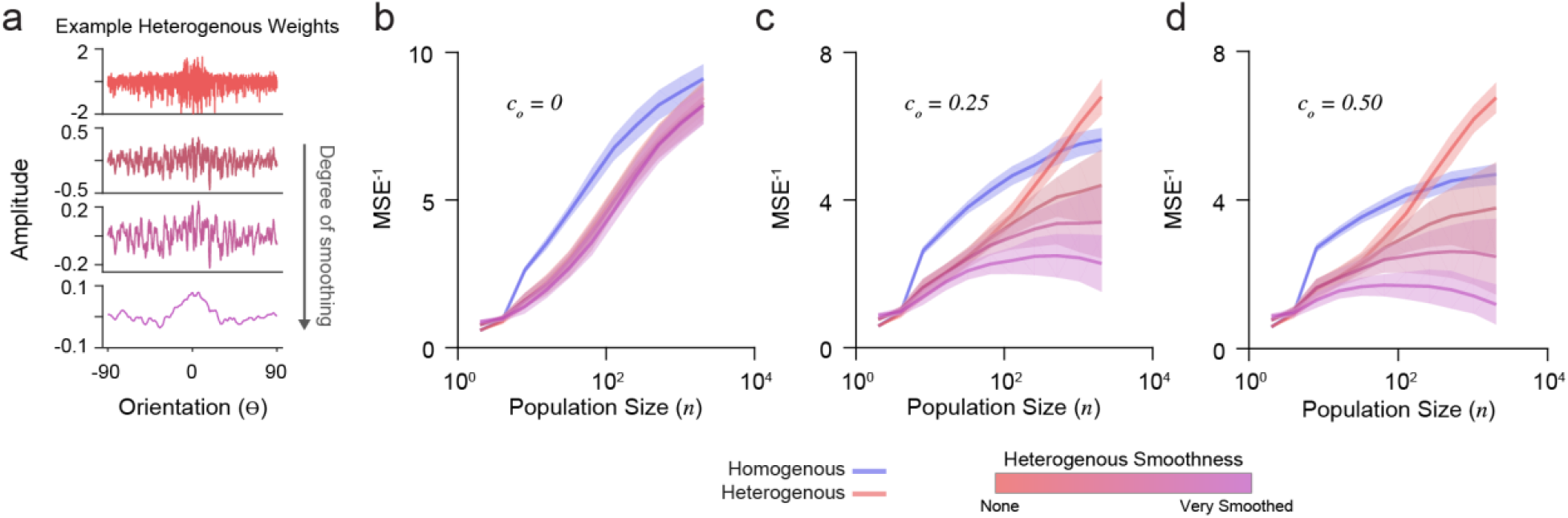
Decoder performance of heterogeneous input populations depends on population size, correlations, and weight diversity. (a) Example weight distribution for a decoder neuron reading out a heterogeneous input population (*top*). Shown are the effects of progressively smoothing weights. Smooth parameters (see Methods) from top to bottom: (0,0), (0.1,1), (0.2,2), (1,10). Ordinate is orientation preference, restricted between −90° and 90° (b) Decoder performance (inverse mean-squared-error) plotted for homogenous and heterogeneous input populations of increasing size. Simulations here include no correlations (*c_o_* = 0). Shading indicates standard error. (c) Same as in (b) for input populations with moderate correlation (*c_o_* = 0.25). (d) Same as in (b) for input populations with stronger correlation (*c_o_* = 0.50).

Without noise-correlations, the accuracy of all decoders increases with population size, with a homogenous *P_IN_* preforming best (Fig. 3b). In the presence of noise-correlations, accuracy saturates for large homogenous *P_IN_* (Fig. 3c-d). As previously shown (Ecker et al., 2011), accuracy for heterogeneous populations with limited-range correlations does not saturate (Fig. 3c-d). However, this depends on weight diversity. Smoothing the weights for heterogeneous *P_IN_* caused saturation and decreased accuracy (Fig. 3c-d), demonstrating that weight amplitude diversity in analytically derived weights distributions are critical for the decoder performance.

### Simulating excitatory weight tuning

In order to compare analytically derived weights with the synaptic inputs onto V1 neurons measured *in vivo*, we generated excitatory synaptic input populations (*P_SYN_*). Under the assumption that synaptic integration is linear, two synapses of equal weight are the same as one synapse with double that weight. This creates a degeneracy where synapse count and size trade off. Because current spine imaging techniques typically capture large synapses and there is no relationship between strength and orientation preference (Scholl et al., 2021), we can assume size is fixed and convert the derived weights into a frequency distribution of ‘synaptic inputs’ (Fig. 4a). The tuning curve for such a synapse is the tuning curve of the input and thus, the synaptic input population, *P_SYN_*, is the input population resampled with probabilities given by the derived weights. An example *P_SYN_* for a single decoder neuron is shown in Figure 4b (drawn from the heterogeneous *P_IN_* in Figure 2). *P_SYN_* in this example displays some specificity in orientation tuning relative to the somatic output, indicated by a larger proportion of simulated synapses with similar orientation preference as the somatic output (0°) of the decoder neuron.

**Figure 4:**
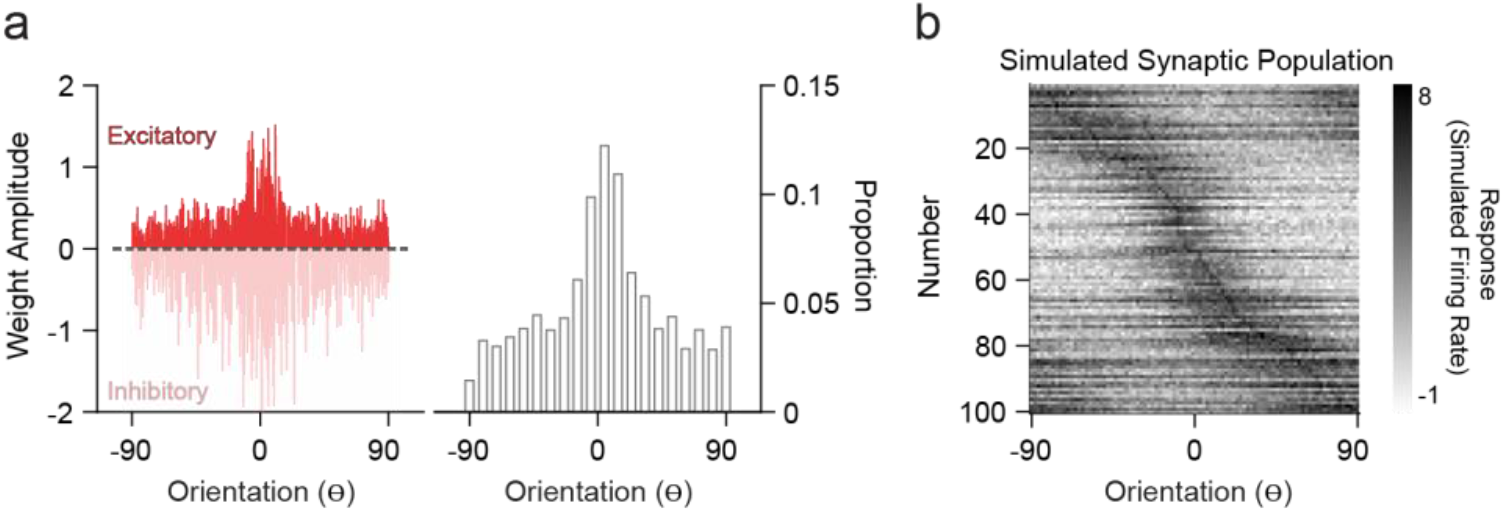
Simulation of synaptic populations from decoder neuron weight distributions. (a) Example weight distribution for a single decoder neuron tuned to 0° (*left*). Ordinates are orientation preference, restricted between −90° and 90° Dashed line separates excitatory (positive) and inhibitory (negative) weights. Excitatory weight distribution over the input population is transformed into a frequency distribution, whereby greater amplitude equates to greater frequency of occurrence (*right*). (b) Example simulated synaptic population (n = 100 spines) from the weight distribution in (a). Shown are the orientation tuning curves of each simulated synapse (normalized).

### Empirical distribution of dendritic spine tuning is consistent with decoding of a heterogeneous input population

We analyzed two-photon calcium recordings from soma and corresponding dendritic spines on individual neurons in ferret V1 during the presentation of oriented drifting gratings (see Methods). While our model draws from a *P_IN_* matched to measurements from macaque V1, the orientation tuning of layer 2/3 neurons in ferret V1, as measured by two-photon cellular imaging, exhibit a similar range in selectivity (Wilson et al., 2017). Visually responsive and isolated dendritic spines (see Methods) typically exhibit diverse orientation tuning relative to the somatic output, although some individual cells show greater overall diversity (Fig. 5b) than others (Fig. 5a). To characterize *P_SYN_* diversity, both for real dendritic spines and simulated inputs, we computed the Pearson correlation coefficient between the tuning curves of individual inputs and the corresponding somatic output (Scholl et al., 2021). For these comparisons, we sampled orientation space in the model spines to match our empirical measurements (22.5 deg increments) and the number of total excitatory inputs recovered for each simulated downstream neuron was set to 100, similar to the average number of visually-responsive spines recorded for each ferret V1 neuron (n = 45, n = 158.9 ± 73.2 spines/cell). Simulations were run 10,000 times, with *N* = 1,000 for *P_IN_* and *c_o_* = 0.20.

**Figure 5:**
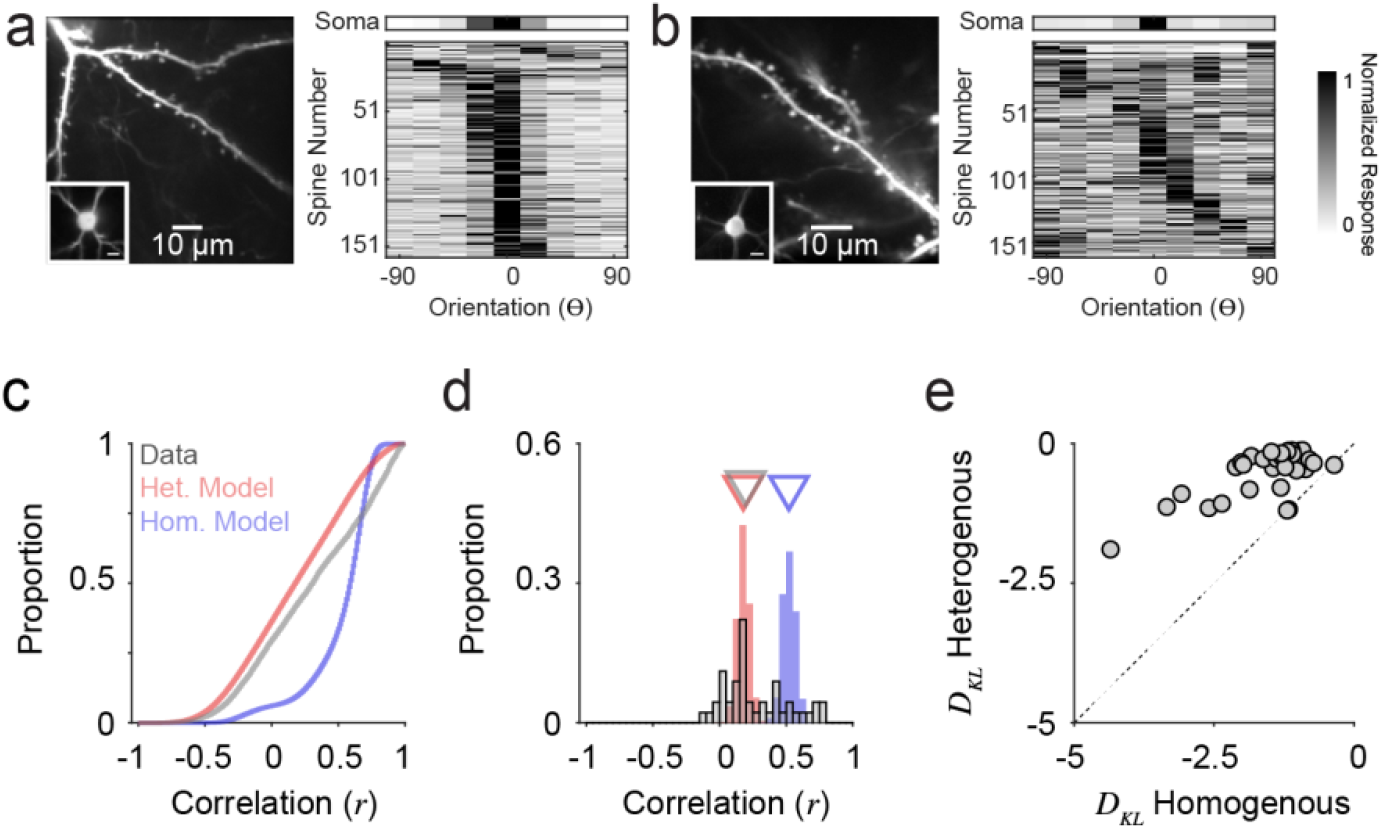
Orientation tuning diversity of dendritic spine populations in ferret V1 match simulations with correlated, heterogeneous input populations. (a) Two-photon standard-deviation projection of example dendrite and spines recorded from a single cell (*left*). Inset: Two-photon standard-deviation projection of corresponding soma. Scale bar is 10 microns. Orientation tuning of soma (*top*) and all visually-responsive dendritic spines from this single cell (n = 159) are shown (*right*). Spine responses are normalized peak ΔF/F. Orientation preferences are shown relative to the somatic preference (aligned to 0°). (b) Same as in (a) for another example cell (n = 162 visually-responsive spines). (c) Cumulative distributions of tuning correlation between individual dendritic spines or simulated synaptic inputs with corresponding somatic tuning or decoder output. Shown are correlations of simulations of homogenous (*blue*) or heterogeneous (*red*) input populations, compared to empirical data (*gray*). (d) Distributions of average tuning correlation between synaptic input and somatic output across measured cells (n = 45). Also shown are distributions of average tuning correlation for simulated cells. Triangles denote median values for each distribution. (e) Comparison of Kullback-Leibler divergence (*D_KL_*) between data and each model type. Each data point represents an individual cell’s population of dendritic spines.

Across all simulated inputs, input-output tuning correlation was higher for homogeneous *P_SYN_* compared to heterogeneous *P_SYN_* (median *r_hom_* = 0.60, median *r_hom_* = 0.18, n = 900,000; Fig. 5c). Tuning correlation between all imaged dendritic spines and soma was low (median *r_cell_* = 0.31, n = 7,151 spines from 45 cells), more closely resembling our model with a heterogenous *P_IN_*. As somatic orientation selectivity (i.e., tuning bandwidth) varies for single cells in ferret V1 (Goris et al., 2015; Wilson et al., 2016), we next examined the average input-output tuning correlation across individual cells (Fig. 5d). Here, the homogeneous model exhibited greater specificity then the heterogeneous model (median *r_hom_* = 0.52; median *r_het_* = 0.18, n = 90,000). For ferret V1 cells, we observed similar spine-soma correlation as the heterogeneous simulation (median *r_cell_* = 0.20, n = 45). Ferret V1 cells were not statistically different from neural decoders with a heterogeneous *P_SYN_* (p = 0.19, Mann-Whitney test), while neural decoders with a homogeneous *P_SYN_* were significantly more correlated with the inputs (p < 0.0001, Mann-Whitney test). A small percentage of imaged cells (17.9%, n = 5/28) had synaptic populations whose mean tuning correlation were within the 95% confidence interval of the homogeneous model distribution. Additionally, some cells had negative average correlations with their spines, which never occurred in the models—potentially indicating nonlinearities between the spines and soma. It is also important to emphasize that both synaptic populations and the heterogeneous model exhibit a positive bias in tuning correlations, illustrating that while inputs are functionally diverse, they are, on average, more similarly tuned to the cell/decoder output.

Given the differences between ferret V1 neurons, we quantified the degree to which synaptic populations on *each* neuron matched tuning correlation distributions from models of homogenous and heterogenous *P_IN_*, by calculating the Kullback-Leibler divergence (*D_KL_*, bin size = 0.05, see Methods). Across our population, imaged neurons more closely resembled simulations with heterogenous, compared to homogenous, *P_IN_* (93.3%, n = 42/45; Fig. 5e) and *D_KL_* from a heterogenous model was consistently larger (p < 0.0001, sign rank Wilcoxon test). This trend held for a range of histogram bin sizes (0.001 – 0.20). Importantly, the models are not fit to data. They are derived entirely from the statistics of the input population, so this correspondence between the heterogenous model and the data results from no free parameters.

In addition to the similarity in input-output tuning correlation, we observed several trends predicted by the heterogeneous model that were evident in synaptic populations imaged *in vivo*. Simulated excitatory inputs correlated with the decoder output were not more selective for orientation (see Methods) (bootstrapped PCA slope = 0.001 ± 0.004 s.e., n = 10,000 simulation runs). For two-photon data, a minuscule, but significant, trend was evident (bootstrapped PCA slope = 0.03 ± 0.1 s.e., n = 7151). So while selective inputs are proposed to provide more information about encoded stimulus variables (Seriès et al., 2004; Shamir and Sompolinsky, 2006; Zavitz and Price, 2019) and unselective (or poorly selective) inputs could convey information through their covariance with selective neurons (Zylberberg, 2017), our model and experimental data suggest co-tuned and orthogonally-tuned inputs exhibit a wide range of tuning selectivity. Response variability (i.e. standard deviation) across trials for simulated excitatory inputs was significantly smaller for ‘null’ orientations (± 90 deg) than at the ‘preferred’ (median = 0.30 and IQR = 0.14, median = 0.38 and IQR = 0.22, respectively; p < 0.001, Wilcoxon ranksum test). This trend was also observed in our two-photon data (null: median = 0.13 and IQR = 0.15; preferred: median = 0.23 and IQR = 0.31, respectively; p < 0.001, Wilcoxon ranksum test). As both modeled and imaged neurons had “null”-tuned excitatory inputs that exhibited less response variability, these inputs may carry useful information about when the preferred stimulus is *not* present.

Taken together, our decoding framework with a realistic (i.e. heterogenous orientation tuning), noisy input populations suggest the collection of orthogonally-tuned excitatory inputs in cortical neurons *in vivo* are not unexpected. Instead, the synaptic architecture of layer 2/3 neurons in ferret visual cortex are likely optimized for the readout of upstream populations tuned to orientation.

## Discussion

We used a population decoding framework (Jazayeri and Movshon, 2006; Kohn et al., 2016; Pouget et al., 2000; Shamir, 2014) to elucidate a possible source of synaptic diversity in functional response properties. We find that even simple decoders exhibit substantial heterogeneity in their weights when the inputs are noisy, correlated neural populations with heterogeneous orientation tuning. We argue that this could naturally explain the heterogeneity in synaptic inputs measured *in vivo* if these cortical neurons are decoding information from upstream input populations. We compared two neural decoders: one with homogenous input (Jazayeri and Movshon, 2006) and one with heterogenous input (Ecker et al., 2011). We show that empirical measurements from dendritic spines recorded within individual cortical neurons in ferret V1 exhibit a similar amount of diversity in orientation tuning as simulated inputs (i.e. excitatory weights) from heterogeneous input populations. It may appear trivial that heterogeneous input populations would produce heterogeneous weights, but it was neither immediately obvious that the weights would not be smooth nor that excitatory weights would be evident for orthogonal orientations. Orthogonally-tuned or non-preferred inputs are often considered to be aberrant; to be pruned away during experience-dependent plasticity or development (Holtmaat and Svoboda, 2009). Our decoding approach suggests these inputs are purposeful and emerge through development as cortical circuits learn the statistics of their inputs (Avitan and Goodhill, 2018). Taken together, our results shed light on synaptic diversity that has been puzzling, suggesting that it is, in fact, expected given known properties of the input population.

We believe our study is a significant step forward in combing population coding theory (Averbeck et al., 2006; Pouget et al., 2000) and functional connectomics (Wilson et al., 2016). The ability to measure receptive field properties and statistics of sensory-driven responses of synapses *in vivo* provides a new testing bed for population codes. The individual neurons which synapses converge on are the real components of what has long been a hypothetical downstream population decoder. While we did not set out to build a computational or biophysical model of a neuron, we believe simplistic approaches such as ours are fruitful for understanding basic principles.

To limit complexity, our decoder did not account for many aspects of cortical networks such as stimulus-dependent correlations or recurrent connections. In the case of stimulus dependent covariance, the optimal decoder is no longer linear, however, that decoder closely resembles a complex cell (Jaini and Burge, 2017; Pagan et al., 2016). Extending a decoding framework to include realistic noise has been used to capture many nonlinear features of neural responses including divisive normalization, gain control, and contrast-dependent temporal dynamics—all features which fall naturally out from a normative framework (Chalk et al., 2017). These more sophisticated approaches may be able to make predictions about the synaptic organization itself, whereby local clusters of synapses act as nonlinear subunits (Ujfalussy et al., 2018).

Our model does not describe a cortical transformation. Instead, to limit complexity, we focused on the propagation of orientation selectivity from one neural population to another, akin to the propagation of basic receptive field properties from V1 to higher-visual areas. Our approach was chosen to provide a starting point for predicting the tuning diversity of synaptic input populations as compared to the tuning output or downstream cells. However, this model could be modified to study the convergence and transformation of cortical inputs. An obvious case study would be complex cells in layer 2/3 V1 (Hubel and Wiesel, 1962; Movshon et al., 1978; Spitzer and Hochstein, 1988), which are thought to integrate across presynaptic cells with similar oriented receptive field with offset spatial subunits to produce polarity invariance. This extension would be better suited for a nonlinear quadratic decoder (Jaini and Burge, 2017; Pagan et al., 2016), rather than the linear one used here. We hope that future studies build upon this modeling framework, exploring quadratic decoders and work towards using richer visual stimuli and neural models (Chalk et al., 2017). We believe this will be critical for gaining insight into how information propagates between cortical areas and from largescale measurements of cortical functional connectivity.

## Materials and Methods

All procedures were performed according to NIH guidelines and approved by the Institutional Animal Care and Use Committee at Max Planck Florida Institute for Neuroscience.

### Derivation for a Bayesian probabilistic decoder

We construct a probabilistic decoder, represented by a population of neurons, that reports or estimates the identity of a stimulus from the spiking response of an input population of neurons. We assume an input population with responses that are a function of the stimulus, *f*(θ*_k_*), plus Gaussian noise, *f*(θ*_k_*), and the covariance (*Q*) is equal for all stimulus conditions (*Q*) such that *Q* = *Q_k_* = *Q_i_*. Then, the posterior distribution can be written as

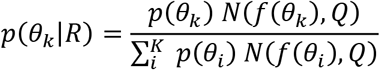

where a multivariate Gaussian is

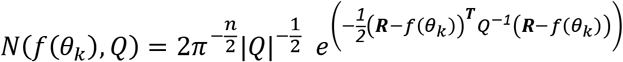

This can be expanded and simplified such that

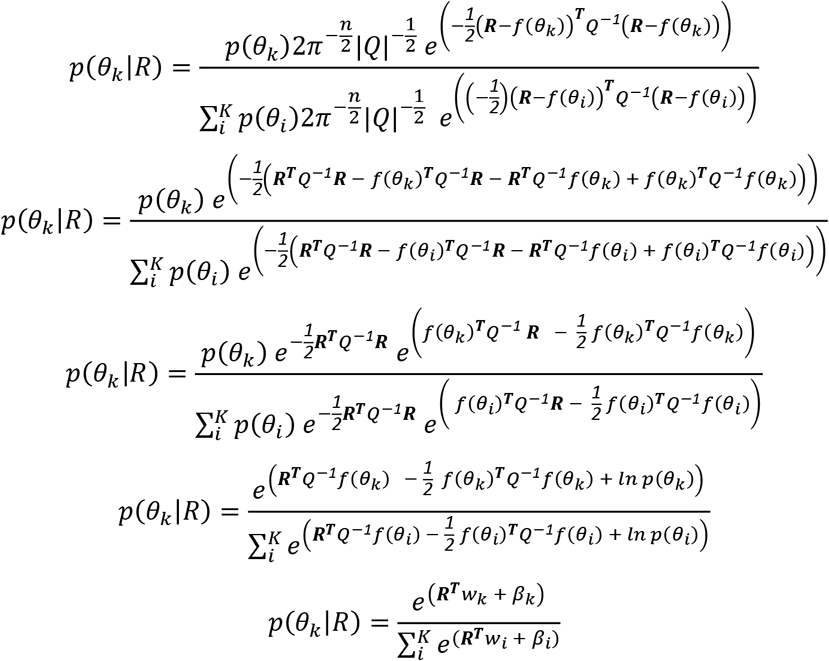

where

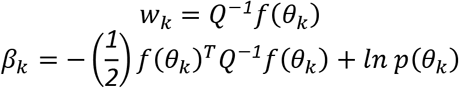

Here, *w* are the weights over *k* for each neuron in the decoder population and *β* is a constant term for each *k*. Importantly, because we assume Gaussian input, with this formulation, *w* and *β* are derived closed form. More generally, *w* and *β* can be estimated numerically using multinomial logistic regression and this form remains optimal for any input population statistics within the exponential family (e.g., Poisson noise).

### Input population model

To generate input populations (*P_IN_*), we simulated *N* neurons responding to a stimulus characterized by orientation (θ*_k_* ∈ [-π/2:π/2*K*:π/2]). The response of each neuron, *r_i_*, depends on a tuning function, *f_i_* (*θ*) and an additive noise term, *ε_i_*, describing trial-to-trial variability. Noise is correlated across the population, generated from a multivariate Gaussian distribution with zero mean and covariance *C*. Orientation tuning functions were defined as:

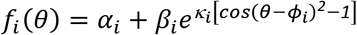

Here, α is the baseline firing rate, β scales the tuned response, κ scales the tuning bandwidth, and ϕ is the orientation preference of each neuron. For homogeneous *P_IN_* all parameters except ϕ were fixed: (α, β, κ) = (0, 5, 4). For heterogenous *P_IN_*, we sampled parameters to match measurements from macaque V1 (Ringach et al., 2002) and our ferret V1 data. Tuning bandwidth was generated by converting half-width at 1/√2 height (γ) values from a lognormal distribution (μ = −1, σ = 0.6):

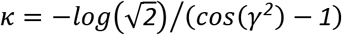

Limited-range correlations were included so neural noise correlation depends on tuning preference difference (Ecker et al., 2011). A correlation matrix, *C*, was specified by the difference between preferred orientations of neurons and the maximum pairwise correlation, *c_o_*:

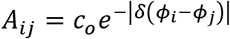

where δ is the circular difference and

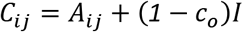

where *I* is the identity matrix of size *N*. We scaled the correlation matrix by the mean firing rate of each neuron to produce Poisson-like noise (Ecker et al., 2011).

Derived weights for a given *P_IN_* were artificially smoothed using the following equation from (Park and Pillow, 2011):

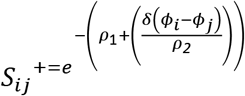

Here, *S*^+^ is the pseudoinverse of *S*, *δ* (ϕ_*i*_ - ϕ_*j*_) is the circular difference between preferred orientations of neurons, ρ_1_ scales the amplitude of smoothing, and ρ_2_ scales functional range of smoothing.

### Population decoder estimation accuracy

Decoding accuracy was calculated with the mean-squared-error of the maximum a posterior probability (*MAP*) estimate across *t* simulated trials of each stimulus (*k*):

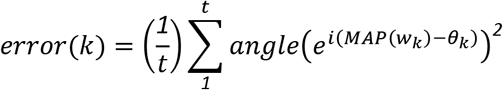

Here, *w_k_* are the weights for a given decoder neuron and θ*_k_* is the true stimulus.

### Viral Injections

Briefly, female ferrets aged P18-23 (Marshall Farms) were anesthetized with isoflurane (delivered in O2). Atropine was administered and a 1:1 mixture of lidocaine and bupivacaine was administered SQ. Animals were maintained at an internal temperature of 37 degrees Celsius. Under sterile surgical conditions, a small craniotomy (0.8 mm diameter) was made over the visual cortex (7-8mm lateral and 2-3mm anterior to lambda). A mixture of diluted AAV1.hSyn.Cre (1:25000 to 1:50000) and AAV1.Syn.FLEX.GCaMP6s (UPenn) was injected (125 - 202.5 nL) through beveled glass micropipettes (10-15 micron outer diameter) at 600, 400, and 200 microns below the pia. Finally, the craniotomy was filled with sterile agarose (Type IIIa, Sigma-Aldrich) and the incision site was sutured.

### Cranial Window

After 3-5 weeks of expression, ferrets were anesthetized with 50mg/kg ketamine and isoflurane. Atropine and bupivacaine were administered, animals were placed on a feedback-controlled heating pad to maintain an internal temperature of 37 degrees Celsius, and intubated to be artificially respirated. Isoflurane was delivered throughout the surgical procedure to maintain a surgical plane of anesthesia. An intravenous cannula was placed to deliver fluids. Tidal CO2, external temperature, and internal temperature were continuously monitored. The scalp was retracted and a custom titanium headplate adhered to the skull (Metabond, Parkell). A craniotomy was performed and the dura retracted to reveal the cortex. One piece of custom cover-glass (3mm diameter, 0.7mm thickness, Warner Instruments) adhered using optical adhesive (71, Norland Products) to custom insert was placed onto the brain to dampen biological motion during imaging. A 1:1 mixture of tropicamide ophthalmic solution (Akorn) and phenylephrine hydrochloride ophthalmic solution (Akorn) was applied to both eyes to dilate the pupils and retract the nictating membranes. Contact lenses were inserted to protect the eyes. Upon completion of the surgical procedure, isoflurane was gradually reduced and pancuronium (2 mg/kg/hr) was delivered IV.

### Visual Stimuli

Visual stimuli were generated using Psychopy (Peirce, 2007). The monitor was placed 25 cm from the animal. Receptive field locations for each cell were hand mapped and the spatial frequency optimized (range: 0.04 to 0.25 cpd). For each soma and dendritic segment, square-wave drifting gratings were presented at 22.5 degree increments (2 second duration, 1 second ISI, 8-10 trials for each field of view).

### Two photon imaging

Two photon imaging was performed on a Bergamo II microscope (Thorlabs) running Scanimage (Pologruto et al., 2003) (Vidrio Technologies) with 940nm dispersion-compensated excitation provided by an Insight DS+ (Spectraphysics). For spine and axon imaging, power after the objective was limited to < 50 mW. Cells were selected for imaging on the basis of their position relative to large blood vessels, responsiveness to visual stimulation, and lack of prolonged calcium transients resulting from over-expression of GCaMP6s. Images were collected at 30 Hz using bidirectional scanning with 512×512 pixel resolution or with custom ROIs (frame rate range: 22 - 50 Hz). Somatic imaging was performed with a resolution of 2.05 - 10.24 pixels/micron. Dendritic spine imaging was performed with a resolution of 6.10 −15.36 pixels/micron.

### Two Photon Imaging Analysis

Imaging data were excluded from analysis if motion along the z-axis was detected. Dendrite images were corrected for in-plane motion via a 2D cross-correlation based approach in MATLAB or using a piecewise non-rigid motion correction algorithm (Pnevmatikakis and Giovannucci, 2017). ROIs (region of interest) were drawn in ImageJ; dendritic ROIs spanned contiguous dendritic segments and spine ROIs were fit with custom software. Mean pixel values for ROIs were computed over the imaging time series and imported into MATLAB (Hiner et al., 2017; Sage et al., 2012). ΔF/F_o_ was computed by computing F_o_ with time-averaged median or percentile filter (10th percentile). For spine signals, we subtracted a scaled version of the dendritic signal to remove back-propagating action potentials as performed previously (Wilson et al., 2016). ΔF/F_o_ traces were synchronized to stimulus triggers sent from Psychopy and collected by Spike2. Spines were included for analysis if the SNR of the preferred response exceeded 2 median absolute deviations above the baseline noise (measured during the blank) and were weakly correlated with the dendritic signal (Spearman’s correlation, r < 0.4). Some spine traces contained negative events after subtraction, so correlations were computed ignoring negative values. We then normalized each spine’s responses so that each spine had equal weight. Preferred orientation for each spine was calculated by fitting responses with a Gaussian tuning curve using lsqcurvefit (Matlab). Tuning selectivity was measured as the vector strength index (*v*) for each neuron’s response:

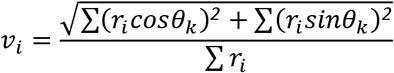

Here *r* is the mean responses over the orientations (θ*_k_*) presented for each spine (*i*). Note, this same index is used to characterize simulated input selectivity.

### Analysis

To compare input tuning (derived synaptic population or measured dendritic spine population) with output tuning (downstream readout or measured somatic tuning) we computed the Pearson Correlation coefficient (Matlab). This correlation was computed on trial-averaged responses across different orientations. For dendritic spines and soma, measured responses across stimulus presentation trials were averaged. For simulated synaptic populations and corresponding downstream readout neuron, we simulated trials by adding noise to each synaptic tuning curve.

## Code Availability

Matlab code to generate input and readout populations used are provided: https://github.com/schollben/SpineProbablisticModel2020

## Acknowledgements

This work was supported by the NIH and Max Planck Florida Institute. Jacob Yates is supported by the K99EY032179. Benjamin Scholl is supported by K99EY031137. We thank Alex Huk, Richard Lange, Sabya Shivkumar, Krishnan Padmanabhan, and Jan Kirchner for helpful comments and discussion.

## Author contributions statement

J.Y and B.S. conceived of the experiment. J.Y. derived analytic solutions, J.Y and B.S. preformed analysis, J.Y and B.S. wrote the manuscript.

